# Collective Sensing in Electric Fish

**DOI:** 10.1101/2023.09.13.557613

**Authors:** Federico Pedraja, Nathaniel B. Sawtell

## Abstract

A number of organisms, including dolphins, bats, and electric fish, possess sophisticated active sensory systems that use self-generated signals (e.g. acoustic or electrical emissions) to probe the environment^1,2^. Studies of active sensing in social groups have typically focused on strategies for minimizing interference from conspecific emissions^2-4^. However, it is well-known from engineering that multiple spatially distributed emitters and receivers can greatly enhance environmental sensing (e.g. multistatic radar and sonar)^5-8^. Here we provide evidence from modeling, neural recordings, and behavioral experiments that the African weakly electric fish *Gnathonemus petersii* utilizes the electrical pulses of conspecifics to extend electrolocation range, discriminate objects, and increase information transmission. These results suggest a novel, collective mode of active sensing in which individual perception is enhanced by the energy emissions of nearby group members.

## Main

From the perspective of an agent actively emitting energy to sense the environment the emissions of other agents could either be a source of interference or provide useful information. While the latter principle is commonly exploited in human engineered sensing systems, ranging from autonomous underwater vehicles to medical imaging devices^5-8^, studies of animals have mainly focused on the former possibility^1,2^. For example, some species of South American electric fish sense their environments using continuous, quasi-sinusoidal electric organ discharges (EODs). During social encounters, fish rapidly shift their EOD frequencies apart to minimize mutual interference. This so-called jamming avoidance response is among the most thoroughly studied vertebrate behaviors^3,4^. However, jamming is unlikely to occur when active emissions are extremely brief, as is the case for the acoustic emissions of dolphins, some bat calls, and the EODs of pulse-type electric fish^1,2,9^. While it has been suggested that some dolphins and bats may “eavesdrop” on the returning echoes of conspecific acoustic emissions^10-14^, evidence that active sensing is actually enhanced in such cases is lacking.

Here we examine the effects of conspecific emissions on electrolocation in the African pulse-type electric fish *Gnathonemus petersii*. Prior studies have shown that *Gnathonemus* use their EODs (∼0.3 ms duration pulses separated by variable intervals of 20-200 ms) to detect, localize and discriminate objects based on their electrical resistance and capacitance as well as for social communication^15-18^. However, the physics of electrolocation suggest the possibility that conspecific EODs could also enhance environmental sensing. Due to the steep fall-off of electrical fields with distance, the spatial range of object detection based on active electrolocation is restricted to a body length or less^1^. As a consequence, electric fields generated by nearby conspecifics could significantly extend object detection range. Furthermore, field and laboratory studies have demonstrated rich social behaviors in African pulse-type species, including schooling^19-21^, cooperative foraging^22^, and pack hunting^23^. Tight coordination of the timing of EODs has also been observed in the form of the so-called “echo response” in which conspecifics transiently synchronize their EODs at brief 12-15 ms intervals^24,25^. In our laboratory, we have observed that groups of *Gnathonemus petersii* adopt closely-spaced spatial configurations in the presence of threat stimuli, including dominant fish of the same species. Video analysis of the positions and relative orientations of groups consisting of two subordinate and one dominant fish revealed that the subordinates often spent minutes at a time less than a body length apart in stereotyped spatial configurations, including conspicuous perpendicular, single-file, and parallel line-up arrangements while also synchronizing their EODs (**Extended Data Fig. 1; Video S1**).

### Conspecific EODs enhance electrolocation range

Motivated by these considerations, we used electric field simulations to test whether object detection could be enhanced by the EODs of nearby conspecifics. A realistic three-dimensional boundary element model was used to calculate the changes in the spatial pattern of EOD-induced current flowing through the fish’s skin due to objects at different locations^26^. Such changes, known as electrical images, are encoded by electroreceptors distributed across the body surface and processed within specialized central nervous system pathways^27,28^. For a 1 cm metal sphere positioned 10 cm away from the head, the fish’s own EOD induces an extremely weak electrical image on the skin (termed here the self-image) (**Fig. 1a**, *left*). In contrast, the EOD of a nearby conspecific in a perpendicular configuration induces a prominent electrical image of the same object (termed here the cons-image) (**Fig. 1a**, *right*). To compare the range of self- (blue) and cons-images (red), we constructed 2D maps of the maximal electrical image amplitude on the skin induced by objects located at various locations relative to the fish. Such maps were constructed for eight different conspecific spatial configurations observed behaviorally (**Fig. 1b-e; Extended Data Figure 2)**. The threshold for object detection, estimated from prior behavioral studies^29,30^, is indicated on the maps by the solid line. Notably, for regions of space near the conspecific, cons-images extend further than self-images, increasing detection range by up to 3- fold (**Fig. 1f)**. There is a simple physical explanation for these results. Self-images decay steeply (as approximately the fourth power of distance) due to the fact that the EOD dissipates both as it travels to the object and again on its return to the fish^1^. In contrast, for objects located near the conspecific, cons-images follow a shorter path directly to the sensory surface of the receiving fish, therefore decaying more gradually (roughly as the square of distance) (**Extended Data Fig. 2a,b**). Funneling of current due to the low resistance path of the conspecific fish body also plays a role in shaping the cons-image (**Extended Data Fig. 2c**). Based on prior modeling, we estimate that the fish’s own EOD-induced current would need to be increased ∼100 fold to obtain an increase in electrolocation range comparable to that provided by a conspecific in a perpendicular orientation^1^.

**Fig 1:**
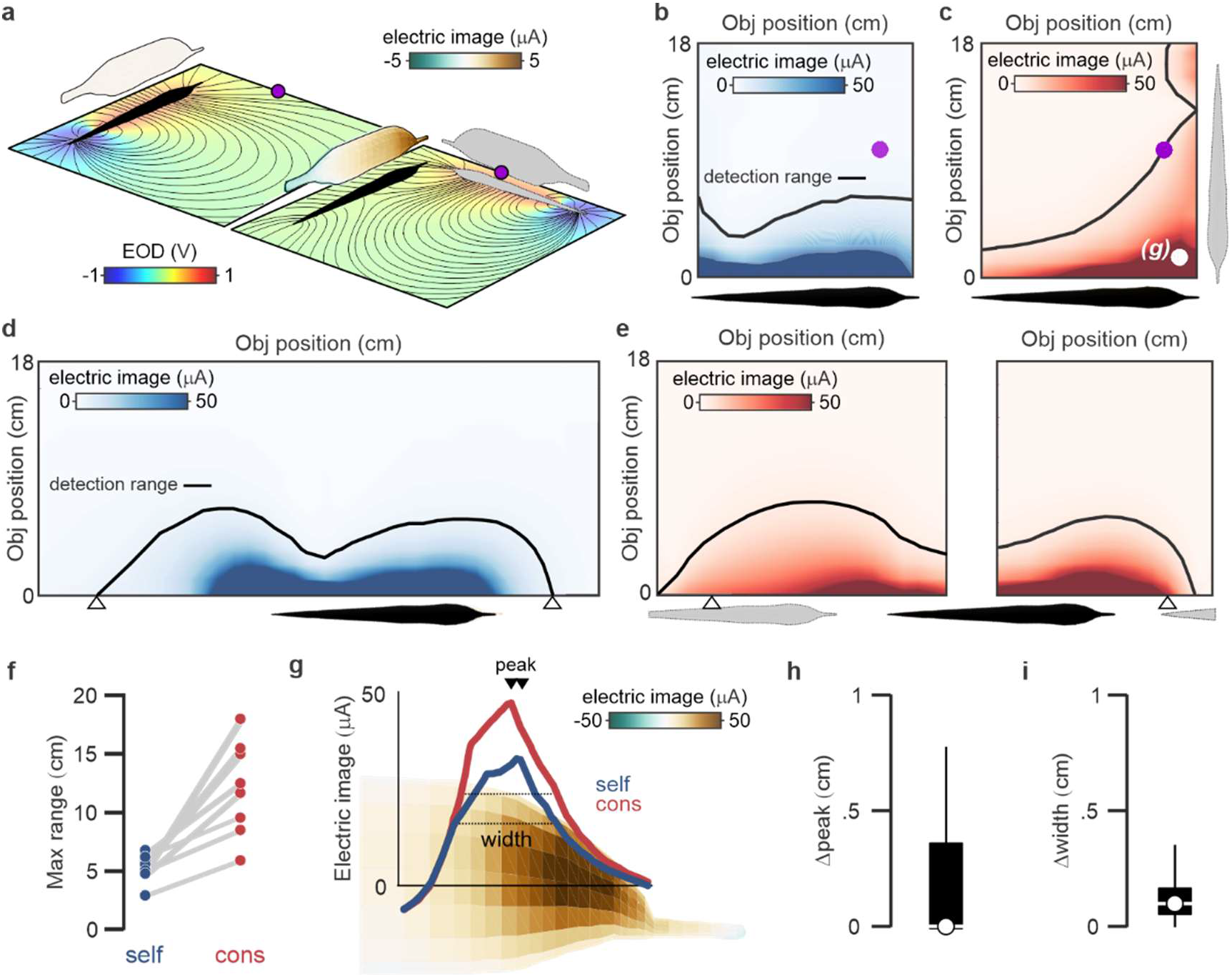
Physical basis for electrolocation based on conspecific EODs. **a**, Heatmaps superimposed on fish contours show boundary element model (BEM) simulations of the electrical images of a 1 cm diameter metal sphere (purple) induced by the fish’s own EOD (self-images) and the EOD of a nearby conspecific shown in gray (cons-images). Electrical images are calculated by subtracting the transcutaneous current in the presence and absence of the object. The plane below the fish contour shows the voltage and electrical field lines induced by the fish’s own EOD (left) and that of the conspecific (right). **b**, Map of the maximal electric image on the skin generated by the fish’s own EOD for a 1 cm metal sphere placed at various distances (y-axis) and locations along the length of the fish (x-axis). Black line indicates the estimated range of object detection. Purple circle indicates the object location simulated in **a. c**, Same display as in **b**, but for electrical images induced by a conspecific EOD. The position of the conspecific is shown by the gray fish contour. White circle indicates the object location simulated in panel **g. d**, Same as in **b** but with an extended x-axis to facilitate comparison with simulations of cons-images due to a single-file configuration of three fish shown in **e**. White arrowheads indicate the limits of the detection based on self-EODs. **e**, Left, cons-image due to the trailing fish (gray) extends electrolocation range to the rear. Right, cons-image due to the leading fish (gray) extends electrolocation range to the front. **f**, Maximal range for self- and cons-images for 8 different conspecific spatial configurations observed behaviorally. **g**, Spatial profiles of self- and cons-images near the head of the fish for the object location and conspecific configuration shown in **c. h,i** Boxplots showing the difference in electric image peak location on the skin and electric image width for self- and cons-images simulated for a range of object positions for 8 different conspecific spatial configurations observed behaviorally. Differences in peak position in **h** reflect object positions near the electric organ in the tail.

The foregoing results raise the important question of whether fish can not only detect, but also localize objects based on cons-images, particularly given that the exact location of conspecifics may not be known. Prior studies have shown that the location of the peak of self-images on the skin is a reliable cue for object location, while the spatial profile (e.g. width) of self-images provides information about object size, shape, and distance^15,31,32^. Further analysis of the simulations described above revealed that the peak locations and spatial profiles of self- and cons-images were typically closely aligned (**Fig. 1g-i; Extended Data Fig. 2d,e**). Hence cons- images could be utilized to localize and characterize objects without explicit knowledge of conspecific location.

### Neural correlates of electrolocation range extension by conspecific EODs

Prior studies of neural responses to cons-EODs have been restricted to brain regions specialized for electrocommunication^17,18^. To identify neural correlates for electrolocation based on cons- EODs, we recorded local field potentials (LFPs) in the hindbrain electrosensory lobe (ELL) where electroreceptors mediating active electrolocation terminate forming a map of the body surface (**Fig. 2a**). After positioning a conspecific nearby the recorded fish in a perpendicular configuration, prominent LFPs were recorded not only in response to the fish’s own EOD (**Fig. 2b**, *blue*) but also to the cons-EOD (**Fig. 2b**, *red*), indicating that cons-EODs strongly activate the ELL. LFP amplitude encodes the strength of electrical images on the skin (up to a distance of around 2 cm) and hence can be used to directly test the range extension for cons-images suggested by the electrical field simulations. To do this, we measured the changes in LFP amplitude for both self- and cons-EODs induced by a plastic cylinder positioned at different distances and locations along the length of the fish. This procedure was repeated for a series of electrode penetrations to obtain LFPs from different locations on the ELL body map (reflecting the strength of electrical images on different parts of the skin). For recordings from anterior regions of the ELL map representing the chin appendage, LFP amplitude was modulated by objects at greater distances for cons-EODs than for self-EODs **(Fig. 2c,d,f; Extended Data Fig. 3)**. In contrast, no such differences were observed for LFP recordings from ELL regions representing the head. These results closely match corresponding boundary element model simulations (**Fig. 2e**), providing neural evidence for an extension of electrolocation range by the EODs of nearby conspecifics.

**Figure 2:**
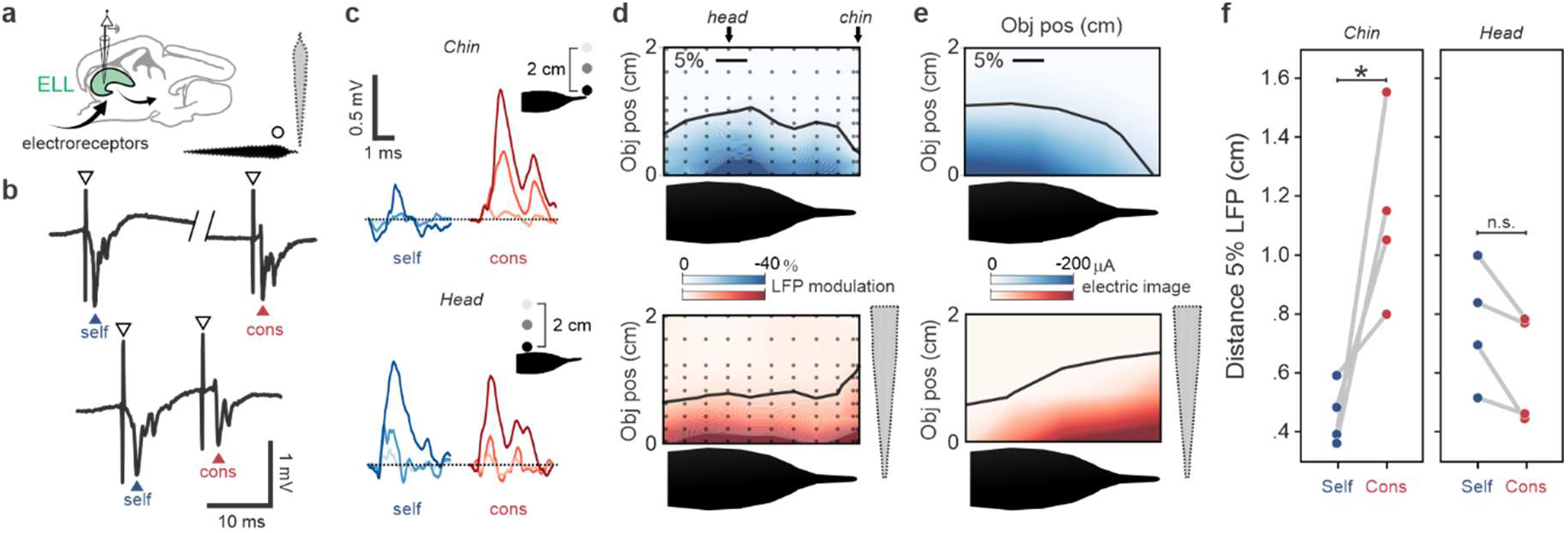
Neural correlates of electrolocation range extension by conspecific EODs. **a**, Schematic of the ELL and spatial configuration of two discharging fish used for LFP recordings. **b**, Two sets of raw LFP traces illustrating responses evoked by self-and cons-EODs. Open arrowheads, EOD artifact. Closed arrowheads, first negative peak of the neural response. **c**, Example traces showing object-induced changes in LFP amplitude evoked by self- (blue) and cons-EODs (red). Responses at three object distances are shown (colored traces). Top, LFP recordings made anteriorly in the chin region of the ELL map. Bottom, traces for recordings in the head region. Note, object induced changes in LFP amplitude are larger for cons- compared to self-EODs in the chin region of the ELL. **d**, Maps of LFP amplitude modulation evoked by a 2 cm plastic rod placed at different locations near the fish (black dots). Each column is separate recording locations on the ELL map matched to the rostro-caudal locations of the object. Top, LFPs evoked by self-EODs. Bottom, LFPs evoked by cons-EODs. Black line indicates the object distance at which LFP amplitude was modulated by 5%. **e**, BEM simulation showing the maximal electrical image amplitude generated by a 2 cm plastic rod placed at different locations near the fish. Top, self-images. Bottom, cons-images. Black line indicates the object distance at which the electrical image was modulated by 5%. Note, range is extended for cons- images when the object is near the chin but not for locations near the head, matching experimental results. **f**, LFPs evoked by cons-EODs are modulated at further distances than those evoked by the fish’s own EOD for object locations near the chin but not for those near the head (Mann Whitney test with sequential Bonferroni significance chin: p = 0.029; head: p = 0.306; n = 4 fish).

### Behavioral evidence for electrolocation range extension by conspecific EODs

Next we sought to provide behavioral evidence that conspecific EODs enhance electrolocation range. Individual fish were housed in a large tank containing two perpendicularly oriented ∼30 cm long alleys, each containing a 2×8 array of “virtual” objects. Each object could be independently switched from an insulator (open circuit) to a conductor via a computer-controlled relay (**Fig. 3a**). A transient acceleration of the fish’s EOD rate following such a switch, known as the novelty response (NR), provided a behavioral indication that the fish detected the change in an object’s electrical properties^33,34^. Two artificial, experimenter-controlled conspecific EOD mimics were placed at the base of each alley near a corner of the tank (**Fig. 3a**, right). Mimics were used in place of live conspecifics to exclude performance increases due to conspecific behavioral responses to objects. The use of two mimics allowed us to test hypothesis from the boundary element modeling regarding the effects of different conspecific configurations on electrolocation range. Although fish were free to move anywhere in the tank, they spent a large majority of the time virtually immobile in close proximity to the conspecific EOD mimics (**Fig. 3b**). Prior studies have shown that fish interact with such mimics similarly to real conspecifics^35,36^. Consistent with this, fish frequently echoed mimic pulses in our setup (**Fig. 3c**).

**Figure 3:**
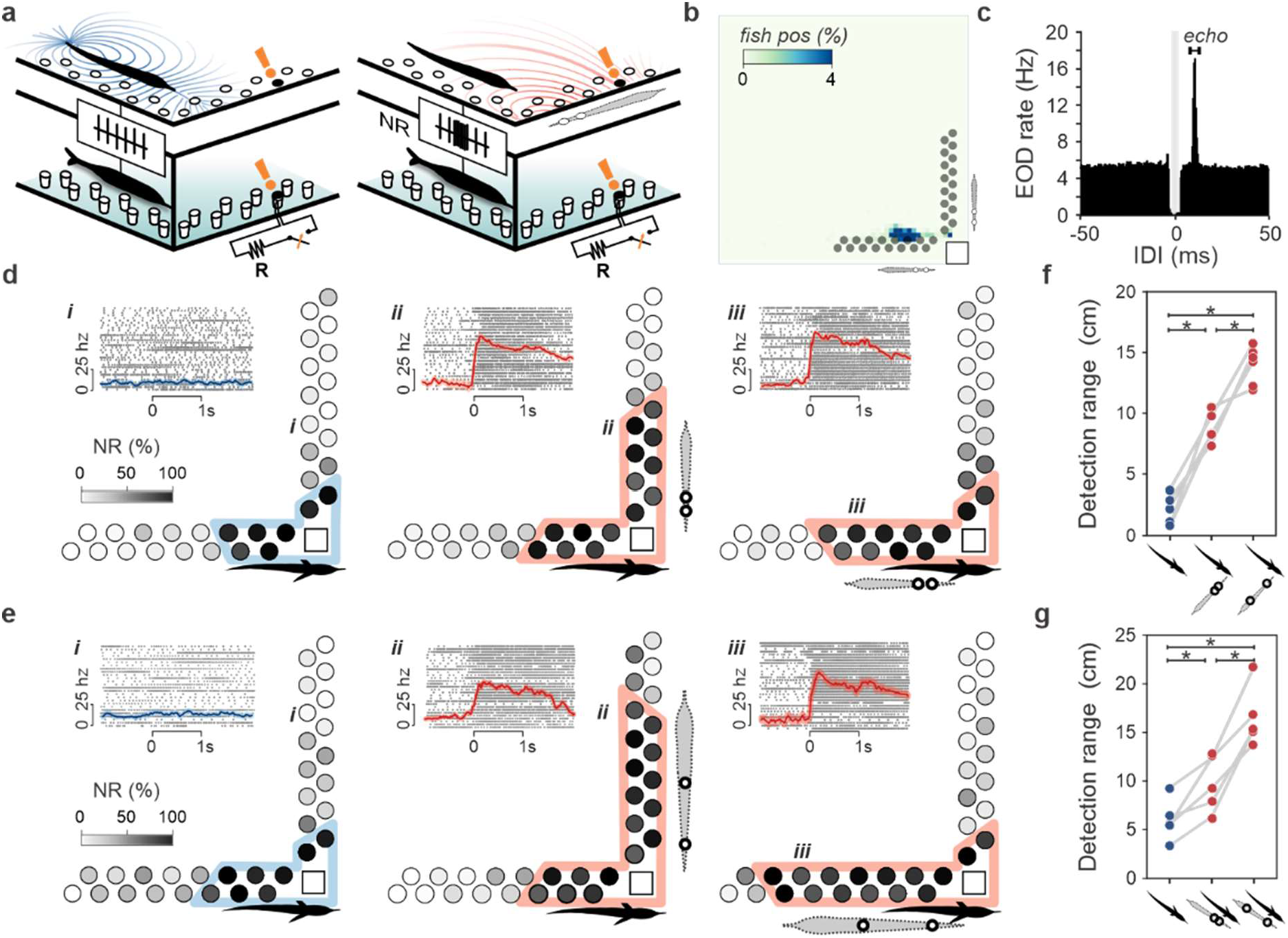
Conspecific EOD mimics increase electrolocation range. **a**, Schematic of the experimental setup. An array of 32 computer-controlled dipole-objects is arranged in an L shape in the corner of the tank. Detection of a change in object properties is measured based on the probability of a transient acceleration of the EOD rate, known as the novelty response (NR), occurring in a brief window following a switch in object resistance (orange). Detection was compared with and without conspecific EOD mimic pulses delivered by one of two pairs of electrodes located in the corners of the tank (gray fish outline). Only one pair is shown for clarity. **b**, Distribution of fish positions in the experimental tank measured over 7 days for a single experiment. **c**, Inter-discharge interval between conspecific EOD mimic and fish revealing prominent echoing of mimic pulses at ∼12 ms. The rate of mimic pulses was ∼10 Hz. EODs could not be reliably detected in the period immediately surrounding the mimic pulse (gray rectangle). **d**, NR probability for each object (grayscale) with mimic pulses off (blue; left) versus on (red; middle, right) for a single fish. Colored contours indicate behavioral detection range (Binomial test: p < 0.05). Insets show raster plots of EOD pulses and average EOD frequency (mean ± SEM) aligned to the time of a switch in resistance of a single object indicated by the roman numeral. Objects that failed to evoke NRs with conspecific EOD mimics off, reliably evoked NRs when the mimic was turned on (compare i and ii). **e**, Same display as **d**, but for a second experiment in the same fish in which the current spread due to the EOD mimic was increased. **f-g**, Summary of changes in detection range across fish and experimental conditions. As expected based on modeling, range increased for both mimic positions and were greatest when the mimic was oriented at a right angle to the fish (n = 5 fish; Mann Whitney test with sequential Bonferroni significance p < 0.05). Increasing mimic current spread increased detection range for both mimic positions. See also **Extended Data Fig 4**

To quantify electrolocation range, a randomly selected object was switched from insulating to conducting for 1 second once every 3 minutes. Such measurements were made in three alternating conditions in which either both mimics were turned off, or one of the two mimics emitted pulses at ∼10 Hz. The fish’s average EOD interval was unchanged in conspecific EOD mimic off versus mimic on conditions (no mimic: 54.6 ± 1.8 ms, mimic: 54.6 ± 2.7 ms, n=5 fish). Data was collected during the light cycle for around one week per fish in order to collect sufficient data for comparing detection range across the three conditions. Object detection range was dramatically increased when the EOD mimics were turned on (**Fig. 3d,f,g, Extended Data Fig. 4**). Distant object locations that seldom or never evoked NRs in the absence of an EOD mimic reliably evoked NRs when the mimic was turned on (**Fig. 3d**, inset). Range was extended in the direction of the field produced by the mimic and was greater when the mimic was oriented perpendicular to the fish (**Fig. 3d,f,g**), closely matching boundary element simulation results (**Extended Data Fig. 5**). In a second round of experiments performed on the same set of fish, we increased the distance between the poles of the EOD mimic (simulating a larger conspecific). As expected based on the model, electrolocation range was extended even further, nearly tripling (**Fig. 3e-g, Extended Data Fig. 4,5**). Continuous video tracking throughout the experiment excluded the possibility that electrolocation range increases were related to changes in fish distance or orientation relative to the objects (**Extended Data Fig. 4**).

### Object discrimination based on conspecific EODs

Weakly electric fish use their own EODs not only to detect objects, but also to discriminate between them based on their size, shape, and electrical properties^15,16^. Therefore, we also tested whether fish can discriminate objects based on a conspecific EOD mimic. Fish received a food reward for entering a chamber behind a resistive object and negative reinforcement (sound or water vibration) for entering a chamber behind a capacitive object. A conspecific EOD mimic was positioned midway between the two objects (**Fig. 4a**). Critically, the electrical properties of the objects were under tight experimental control (see **Methods**; **Fig. 4b**). Testing consisted of two main trial type designed to isolate discrimination performance based on the fish’s own EOD versus that of mimic pulses. In *self* trials, objects differed in their electrical properties throughout the trial and no mimic was delivered (**Fig. 4c**), hence discrimination relied on electrical information extracted from the fish’s own EOD. In *cons* trials, a conspecific mimic pulse was delivered 12 ms after online detection of the fish’s own EOD and the objects differed in their electrical properties only during a brief time window (∼10 ms) bracketing the mimic pulse (**Fig. 4c**). Hence on *cons* trials, the fish’s own EOD contained no information about object properties and discrimination could only be performed based on information extracted from the conspecific EOD mimic. Fish performed above chance on *self* trials from the start of testing, presumably due to a previously reported innate avoidance of large capacitive objects^37^. Remarkably, performance on *cons* trials was indistinguishable from *self* trials, demonstrating that *Gnathonemus* discriminate the electrical properties of objects based on exogenous EODs even when the fish’s own concurrently emitted pulses provide no information (**Fig. 4d**).

**Figure 4:**
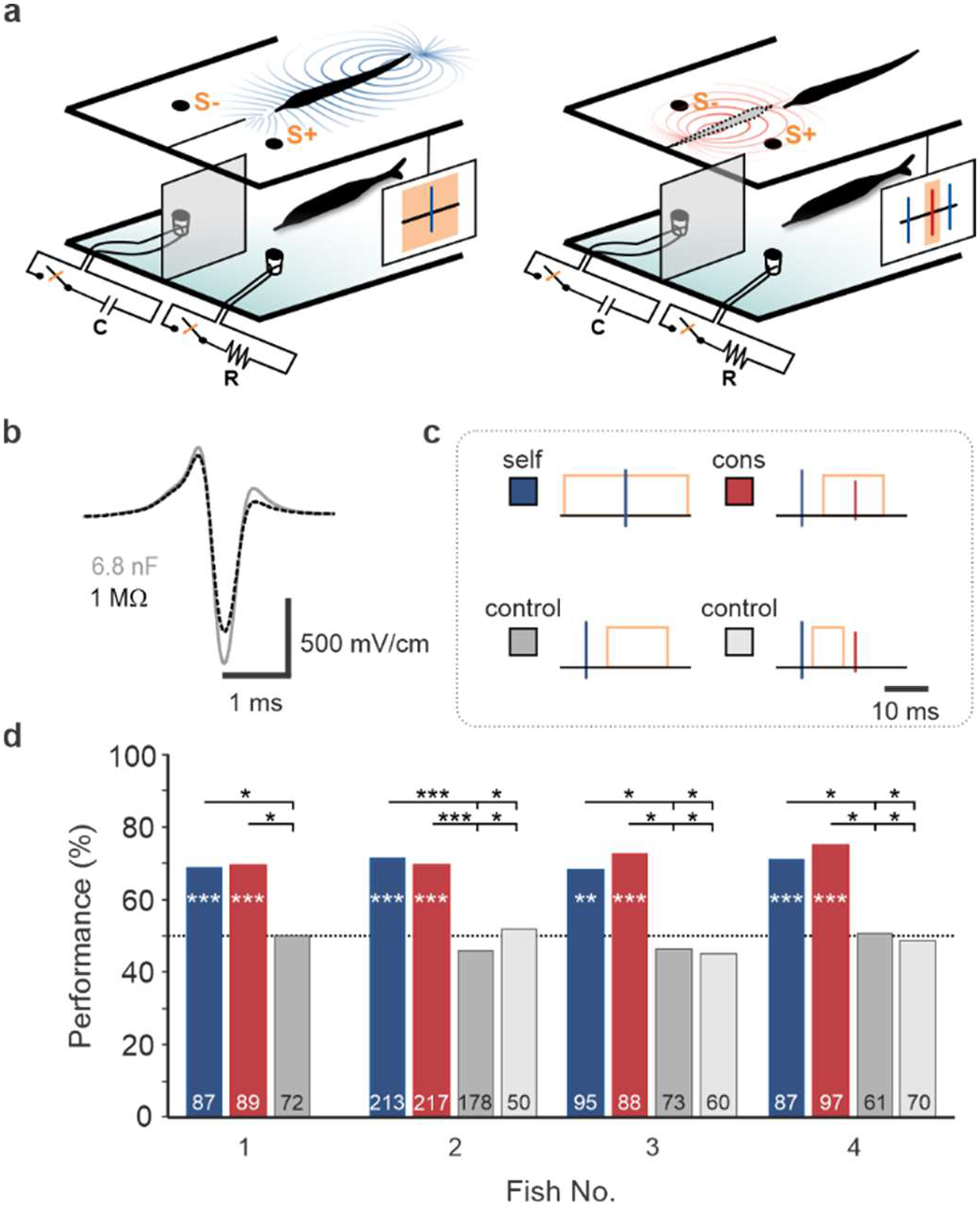
Object discrimination based on conspecific EOD mimics. **a**, Schematic of the experimental setup. Fish discriminated between two objects (S+ and S-) to obtain a food reward. The electrical properties of the objects were controlled by an electronic switch, such that the objects differed in their electrical impedances when the switch was closed (orange rectangle) but were identical when it was open. Discrimination performance was compared with and without conspecific EOD mimic pulses delivered by a pair of electrodes located in from of the object compartment (gray fish outline). **b**, Modulation of the conspecific EOD mimic induced by the two objects (S+, black dashed; S-, gray). **c**, Schematics illustrating the different trial types used to test discrimination. Blue lines indicate the fish’s own EOD, red lines indicate mimic pulses, and the orange rectangles indicate the time period within the trial when object electrical properties differed. **d**, Overall discrimination performance for four fish. The total number of trials per condition per fish is indicated. Binomial tests were conducted to determine if the performances significantly deviated from chance level (*p ≤ 0.05; **p ≤ 0.01; ***p ≤ 0.001). Fisher tests were used to compare the performances between the different conditions for each fish (*p ≤ 0.05; **p ≤ 0.01; ***p ≤ 0.001).

### Conspecific EODs increase electrosensory information transmission rates

Corollary discharge signals associated with the fish’s own EOD motor command are prominent in the ELL and have been hypothesized to enhance, or “gate-in,” electrosensory input related to the fish’s own EOD^27,38^. Strict gating would preclude the transmission of cons-images to higher processing stages. To examine this, we recorded extracellularly from putative output cells of the ELL in an immobilized preparation in which the fish’s own EOD is blocked but the motor command to discharge the electric organ and the associated corollary discharge signals remain intact. Self-images were simulated by varying the amplitude of an artificial EOD pulse delivered at the delay at which the fish’s own pulse normally occurs (4.5 ms), while cons-images were simulated by varying the amplitude of pulses delivered at other delays. Increases and decreases in pulse amplitude simulate changes due to conducting and nonconducting objects, respectively. Importantly, we found that ELL output cells encoded the amplitude of self- and cons-pulses similarly (**Fig. 5a,b; Extended Data Fig. 6a,b**), arguing against strict gating of ELL output by corollary discharge.

**Figure 5:**
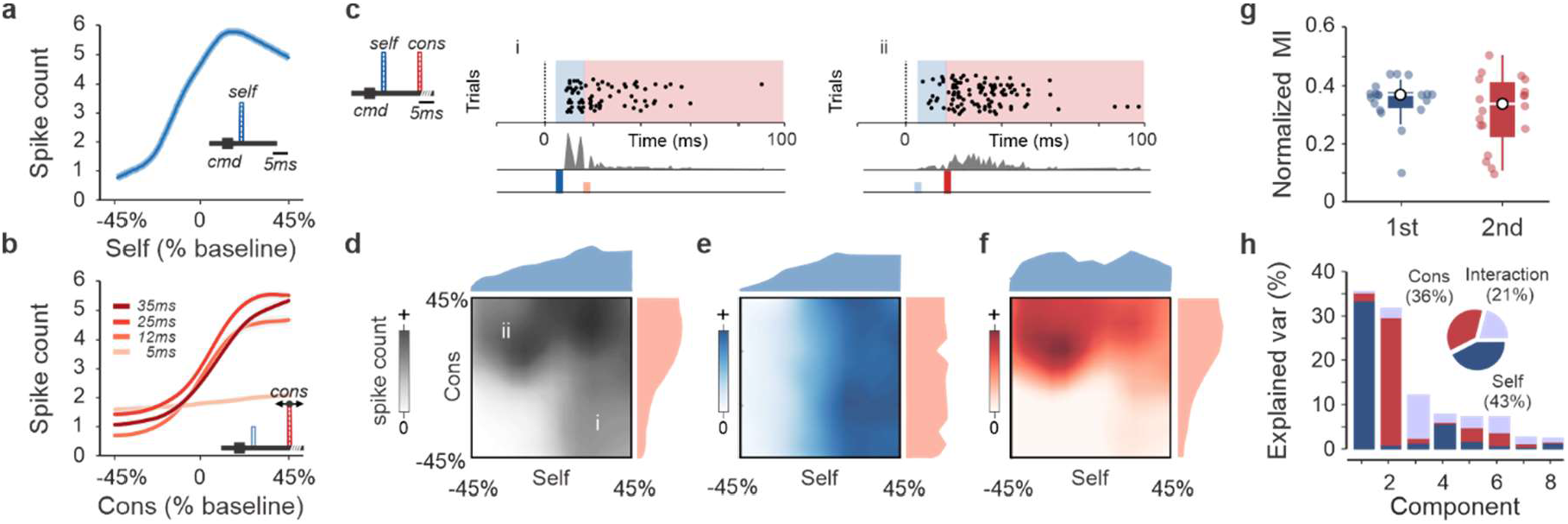
Cons-images increase electrosensory information transmission rates. **a**, Response of an ELL output cell to modulations of the amplitude (3% steps) of a self-EOD pulse (i.e. an artificial pulse EOD locked to the fish’s spontaneously emitted EOD motor command at a 4.5 ms delay corresponding to that of the naturally occurring EOD). **b**, Response of the same cell to modulations of the amplitude of cons-EOD pulses delivered across a range of delays. The low sensitivity at the 5 ms delay is expected due to partial refractoriness in electroreceptors^39^. **c**, Response of an example ELL output cell to independent modulations of self- and cons-pulses (12 ms delay). Rasters and peri- stimulus time histograms illustrating responses to self- (blue) and cons-pulses (red) of different amplitudes (i and ii). Shaded rectangles indicate time windows used for quantifying responses to self- (blue) and cons-pulses (red). **d**, Same unit as in **c**. Grayscale heatmap indicates the total spike count in 100 ms window following the EOD motor command. **e,f**, Responses of the same unit calculated in separate time windows following self- and cons-pulses. **g**, Information about stimulus amplitude for self- and con-pulse amplitudes conveyed by individual ELL output neurons calculated in separate time windows following self- and cons-pulses (n = 19). The normalized mutual information (MI) between both self- and con-pulse amplitudes and firing rate was significantly different from a distribution of MI obtained through random permutations of stimulus amplitude (Control: 0.03 ± 0.02, Mann-Whitney test with sequential Bonferroni significance, p = 1.14e^−7^ for self- and p = 1.27e^−7^ for cons; n = 19). **h**, Demixed principal component analysis for the recorded population of ELL neurons (n = 19). The variance in neuronal responses was mainly explained by components corresponding to self- and cons- pulses with a smaller contribution from mixtures of these two factors.

Finally, we tested whether ELL output neurons are capable of transmitting both self- and cons- image information in the context of the extremely brief (∼12 ms) intervals between self- and cons-pulses characteristic of the echo response(**Fig. 5c; Extended Data Figure 6)**^24,25^. Analyzing spike counts in separate time windows following self- and cons-pulses showed that ELL output neurons faithfully transmit information about the amplitude of both self- and cons-pulses under such conditions despite their close temporal proximity (**Fig. 5d-g**). A population level analysis also confirmed that the output of the ELL provides largely separate representations of the amplitude of self- and cons-pulses (**Fig. 5h**). Given that intervals between self-EODs typically range from 50-200 ms^36,40^, these results strongly support the possibility that social groups enhance electrosensory information transmission rate. Separate decoding windows for self- and cons-image information could be implemented biologically by electric organ corollary discharge signals, which are known to be prominent at higher stages of electrosensory processing in the midbrain, telencephalon, and cerebellum^41-43^.

## Discussion

Results presented here suggest a novel, collective mode of active sensing in *Gnathonemus peterssii* in which environmental perception is enhanced by the energy emissions of nearby conspecifics. In previously described forms of collective sensing (e.g. in fish schools or bird flocks), information is transmitted between group members through behavioral responses to external stimuli^44^. In contrast, sensory information contained in the active emissions of electric fish is instantly and simultaneously available to all group members at no additional cost to the individual emitter in terms of energy expenditure or conspicuousness to predators. Whether similar strategies are employed by other active sensing animals remains an open question^2^, however, aspects of the physics of active electrolocation may be particularly favorable for collective sensing. The large fractional increase in electrolocation range conferred by conspecific groups derives from the short range of active electrolocation (typically less than a body length compared to tens or hundreds of meters for echolocation)^1^. Furthermore, object localization based on conspecific emissions is likely simpler for active electrocation than for echolocation. In the latter case, acoustic emissions must be matched with their corresponding echoes and additional information regarding conspecific location is likely required^11^.

The ecological functions, social organization, and neural computations related to collective sensing in electric fish provide fascinating topics for future research relevant to artificial sensing and autonomous robots as well as neuroscience and sensory biology. While sheltering during the day, the extended range and increased temporal sampling afforded by conspecific groups could serve as an early warning system for detecting predators or conspecific aggressors. Electrolocation based on conspecific EODs may also facilitate cooperative foraging and groups hunting behaviors in which swim fish in closely-spaced formations^20,22,24,45,46^. Cons-images could also enhance object discrimination by providing different views of the same scene. Consistent with this possibility, using our boundary element model simulation it was easy to find scenarios in which two different objects (e.g. a cone versus a cylinder) generated virtually identical self-images but could readily be discriminated based on cons-images (**Extended Data Fig. 7**). Though previously discussed in relation to social communication, the echo response may serve to coordinate collective electrosensing by enforcing rapid turn-taking in which fish alternate discharges. Consistent with such a role, the threshold for the echo response corresponds to that of electroreceptors mediating active electrolocation, rather than the far more sensitive receptors mediating electrocommunication^24,25,45^. This would ensure that turn-taking only occurs between nearby fish who stand to gain additional environmental information from one another’s pulses. Finally, prior work has shown that the cerebellum-like circuitry of the mormyrid ELL implements an internal model that predicts and cancels self-generated electrosensory input related to the fish’s own movements^34,47^. The massively enlarged cerebellum of *Gnathonemus* projects heavily to electrosensory processing structures and hence could potentially support collective sensing by relaying predictions of the electrosensory consequences of conspecific behavior^48-50^.

## Methods

### Experimental Model and Subject Details

Adult male and female Mormyrid fish (11-19 cm in length) of the species *Gnathonemus petersii* were used for experiments. Fish were housed in 60-gallon tanks in groups of 3-20. Water conductivity was maintained between 60-100 μS both in the fish’s home tanks and during experiments. All experiments performed in this study adhere to the American Physiological Society’s Guiding Principles in the Care and Use of Animals and were approved by the Institutional Animal Care and Use Committee of Columbia University.

### Tracking group behavior

Behavioral monitoring was performed in a 60 × 60 cm tank containing an opaque acrylic shelter in one corner and a 12 hr/12 hr light-dark cycle. Groups consisting of one large and two smaller fish were filmed continuously at 150 fps for 3 days under infra-red illumination (USB 3.0 FLIR Chameleon Camera, cm3-U3-13Y3M-CS). Groups of two were not used due to extreme aggression observed under these circumstances^51^. EODs were recorded using a pair of chlorided silver wires and digitized at 30 kHz. The fish were fed once a day at the end of the light cycle. DeepLabCut was used for automated tracking of 5 features (tip of chin appendage, mouth, two points on trunk, and tail) for each of the two smaller fish^52^. 640 manually labelled frames were used for training. Only video frames with tracking confidence levels exceeding 0.95 were included in subsequent analyses.

### Analysis of group behavior

For each video frame, the five tracked points from each fish were used to extract conspecific spatial configurations including: the angle between the head-to-tail axis of the two fish, the minimum Euclidean distance between tracked points belonging to the two fish, and the Euclidean distances between the heads and tails of the two fish. We used clustering analysis to characterize the relationship between the postures of the two fish. Hierarchical clustering with a squared Euclidean distance and Ward’s criterion was used to estimate the number of clusters. K-means clustering was performed in MATLAB for values of k ranging from 2-50. A squared Euclidean distance and preclustering were used to achieve consistent results^53^. The stability and quality of identified clusters was evaluated as described previously^54^. For subsequent boundary element modeling we focused on 7 *social* clusters (out of a total of 11) characterized by a minimum Euclidean distance between the two smaller fish of less than one body length. A hidden Markov Model (HMM) analysis of the transition probabilities between clusters was used to estimate the average continuous time spent within each cluster.

### Boundary element modeling of electrical images

Electric images were computed with software originally developed by Rother and validated in subsequent studies^26,55^. This model has two parts, a geometric reconstruction of the fish’s body and a calculation of the transcutaneous field by solving the Poisson equation for the fish boundary using the Boundary Element Method. Briefly, this method determines the boundary electrical distributions solving a linear system of M · N equations for M poles and N nodes, with the unknown variables being the trans-epithelial current density and potential at each node. The trans-epithelial current density and potential is calculated for each node and linearly interpolated for the triangles defined by the nodes, forming the geometry of fish and objects. Electric images were calculated as the difference between amplitude of positive EOD peak in presence and absence of the object for both self- and cons- EODs.

### Surgery

Fish were anesthetized (MS:222, 1:25,000) and held against a foam pad. Aerated water with MS:222 was continually passed over the fish’s gills for respiration throughout the surgery. Skin on the dorsal surface of the head was removed and a long-lasting local anesthetic (0.75% Bupivacaine) was applied to the wound margins. A 2 mm diameter hole was then drilled in the posterior portion of the skull overlying the medial zone of ELL.

### Electrophysiology

#### LFP recordings in discharging fish

Following the surgery, respiration was switched to tank water containing the sedative Metomidate hydrochloride (Aquacalm; 50 μl/l). Although EOD rate is slowed to ∼1 Hz by sedation, EOD amplitude and waveform and EOD-evoked LFPs in the ELL are minimally altered^56^. LFPs were recorded in medial zone of the ELL with a glass microelectrode positioned in the granular layer. The location of the receptive field on the skin was determined based on LFP responses to brief pulses delivered with a Ag-AgCl dipole electrode held close to the fish’s skin. Once the location of the receptive field was identified on the skin, a 2 cm plastic cylinder was placed at different distances (from 0 to 2 cm in steps of 2-4 mm).

#### Single-unit recordings in paralyzed fish

Surgery was conducted as detailed above to expose ELL for recording. Gallamine triethiodide (Flaxedil) was given at the end of the surgery (∼20 μg/cm of body length) and the anesthetic was removed. Aerated water was passed over the fish’s gills for respiration. Paralysis blocks the effect of electromotoneurons on the electric organ, preventing the EOD, but the motor command signal that would normally elicit an EOD continues to be emitted by the electromotoneurons at ∼5 Hz. The EOD motor command signal was recorded with an Ag-AgCl electrode placed over the electric organ. The command signal lasts about 3 ms and consists of a small negative wave followed by three larger biphasic waves. Onset of EOD command was recorded as the negative peak of the first large biphasic wave in the command signal. Extracellular single-unit recordings were made using glass microelectrodes (2-10 MΩ) filled with 2M NaCl, as described previously^57^. Recording locations within medial zone of the ELL were established using characteristic field potentials evoked by the EOD command^58^. Recordings were restricted to units with receptive fields on the face. Putative output cells were identified based on characteristic responses to electrosensory stimuli and the EOD motor command established in prior studies using *in vivo* intracellular recordings, biocytin labeling, and post hoc histology^59,60^. Both E-type units (corresponding to large fusiform cells) and I-type units (corresponding to large ganglion cells) were analyzed. Electrosensory stimuli were delivered using a chlorided silver dipole electrode placed ∼1 mm from the skin in the region of the receptive field. Data were recorded in Spike2 software (Cambridge Electronic Design) and digitized at 20 kHz.

### Information Content

To determine the information that the spiking response r contains about the stimulus amplitude s, the normalized mutual information (U) was estimated for each time window (1st and 2nd) as:

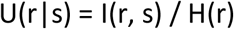

Where I represents mutual information, and H represents information entropy.

Note that 0 <= U(r|s) <= 1. When U(r|s) = 0, the spiking is entirely independent of the stimulus, whereas U(r|s) = 1 indicates that the firing is entirely predictable based on the stimulus amplitude.

### Activity modes from demixed PCA

Demixed PCA was applied to the recorded cells (19 E- and I-cells recorded from 6 fish) using the dPCA package available here (https://github.com/machenslab/dPCA) with the default parameter settings^61^. The input to the dPCA was an n · s1 · s2 · t · k matrix where each entry was the spike rates of individual neurons in individual trials. n corresponds to neurons, s1 corresponds in this case to self-EODs, while s2 corresponds to cons-EODs, t corresponds to time steps (1 ms) and k corresponds to individual trials.

### Object discrimination behavior

The experimental tank was partitioned into a living area (30 × 20 cm; water level, 20 cm) and an experimental area (9 × 12 cm) that was separated by a plastic gate. Fish were habituated to the tank for 1 week prior to the start of the experiment. For testing, a custom-made plastic chamber divided into two compartments was placed in front of the plastic gate. Each compartment contained a dipole object consisting of two carbon poles (diameter: 10 mm; distance between poles: 9 mm). The carbon poles were connected to a switch that allowed the electrical properties of the objects to be controlled. When the switch was open the electrical properties of the two objects were the same. However, when the switch was closed one object (the positive stimulus [S+]), was a 1 MΩ resistor, while the other (the negative stimulus [S-]) was a capacitor of 6.8 nF. For the resistive dipole object, an additional large capacitor (10 μF) was placed in series between the poles of that object (condenser coupling). This large capacitance was far above the upper threshold of all fish, and its impedance for the fish’s EOD was near zero, preventing direct currents from serving as potential additional cues for the fish. Hence object discrimination based on electrolocation was only possible when the switch was closed. The location of the S+ and S- were chosen pseudo-randomly on each trial. EOD mimics were delivered by a carbon dipole electrode placed midway between the objects in front of the divider. The mimic was a pre-recorded conspecific EOD waveform output from a signal generator (Siglent SDG6000X) to a stimulus isolation unit (a-m systems model 2200; peak to peak amplitude: 0.40 mA). Experiments were performed during the light cycle and recorded at 150 fps using a USB 3.0 FLIR Chameleon Camera. The EODs of the fish and mimics were recorded (digitized at 30 kHz, Open Ephys board). Triggering of mimic pulses on self-EODs was performed using analog hardware (Tektronix 26G3). Trials began with opening the gate and ended with the fish swimming back to the living area. A choice was scored when the fish crossed the entrance to the plastic chamber. Correct choices were rewarded with food (a live blackworm) while incorrect choices were punished by using sound or water vibration. Training was conducted five days a week, with each session consisting of 39-45 trials. In each session, three out of four testing conditions were presented in random blocks as schematized in Fig. 3c.

### Behavioral measurements of electrolocation range using the novelty response

Individual fish were housed in a 60 × 60 cm tank (12 hr/12 hr light-dark cycle) with two perpendicularly oriented ∼30 cm long alleys each containing a 2×8 array of “virtual” objects. Each object could be independently switched from an insulator (the baseline state) to a conductor via a computer-controlled homemade EIB with 32 relays connected to a microcontroller (Arduino DUE). A randomly selected object was switched to a conductor for 1 second once every 3 minutes for 20 hours per day for a period of ∼7 days. Two pairs of chlorided silver electrodes located at the base of each alley in one corner of the tank were used to mimic conspecific EODs. Dipole spacing was 4 cm for experiment 1 and 16 cm for experiment 2. The EOD mimic was a biphasic square pulse (0.33 ms width) with a peak-to-peak amplitude of 0.35 mA delivered at 9-11 Hz using a stimulus isolation unit (StimJim) controlled by Bonsai. The experiment consisted of three alternating 20-minute blocks (no mimic, mimic 1, and mimic 2). Transient increases in EOD rate, known as novelty responses (NRs), were detected based on the z-transformed instantaneous EOD rate calculated in 20-minute blocks. A NR was counted when the EOD frequency exceeded a z-value of 1.5 for a minimum of four consecutive EODs. Detection was evaluated for each object based on the fraction of switches that evoked a NR. The experiments were video recorded continuously at 150 fps using a USB 3.0 FLIR Chameleon camera and the EODs of fish and mimics were recorded and digitized at 30 kHz (OpenEphys).

### Quantification and Statistical Analysis

Data were analyzed off-line using Spike2 (Cambridge Electronic Design) and custom Matlab code (Mathworks, Natick, MA). Non-parametric tests were used for testing statistical significance. Differences were considered significant at *P* < 0.05.

## Data and code availability

All data and code will be made available prior to publication on a public repository.

## Acknowledgements

This work was supported by a grant from the NIH (NS075023) to N.B.S. We thank LF Abbott, D. Turcu, and A. Wallach for valuable discussions and comments on the manuscript.

Zuckerman Mind Brain Behavior Institute, Department of Neuroscience, Columbia University, New York, NY 10027

Federico Pedraja and Nathaniel B. Sawtell

## Contributions

N.B.S. and F.P. conceived of the project, designed and performed the experiments and wrote the manuscript. F.P. analyzed the data and performed the modeling.

## Competing financial interests

The authors declare no competing financial interests.

## Notes

### Competing Interest Statement

The authors have declared no competing interest.

## Main References

1 Nelson, M. E. & MacIver, M. A. Sensory acquisition in active sensing systems. J Comp Physiol A Neuroethol. Sens. Neural Behav. Physiol 192, 573–586 (2006).

2 Jones, T. K., Allen, K. M. & Moss, C. F. Communication with self, friends and foes in activesensing animals. J Exp Biol 224, doi:10.1242/jeb.242637 (2021).

3 Heiligenberg, W. Neural nets in electric fish. (MIT press, 1991).

4 Rose, G. J. Insights into neural mechanisms and evolution of behaviour from electric fish. Nat Rev Neurosci 5, 943–951, doi:10.1038/nrn1558 (2004).

5 Braca, P., Goldhahn, R., Ferri, G. & LePage, K. D. Distributed information fusion in multistatic sensor networks for underwater surveillance. IEEE Sensors Journal 16, 4003–4014 (2015).

6 Chernyak, V. S. Fundamentals of multisite radar systems: multistatic radars and multistatic radar systems. (CRC press, 1998).

7 Boyer, F., Lebastard, V., Chevallereau, C., Mintchev, S. & Stefanini, C. Underwater navigation based on passive electric sense: New perspectives for underwater docking. The International Journal of Robotics Research 34, 1228–1250 (2015).

8 Flohr, T. G. et al. Multi–detector row CT systems and image-reconstruction techniques. Radiology 235, 756–773 (2005).

9 Heiligenberg, W. Electrolocation jamming avoidance in the mormyrid fish Brienomyrus. J. Comp. Physiol 109, 357–372 (1976).

10 Gregg, J. D., Dudzinski, K. M. & Smith, H. V. Do dolphins eavesdrop on the echolocation signals of conspecifics? International Journal of Comparative Psychology 20 (2007).

11 Kuc, R. Object localization from acoustic emissions produced by other sonars (L). The Journal of the Acoustical Society of America 112, 1753–1755 (2002).

12 Dechmann, D. K. et al. Experimental evidence for group hunting via eavesdropping in echolocating bats. Proc Biol Sci 276, 2721–2728, doi:10.1098/rspb.2009.0473 (2009).

13 Götz, T., Verfuß, U. K. & Schnitzler, H.-U. ‘Eavesdropping’in wild rough-toothed dolphins (Steno bredanensis)? Biology letters 2, 5–7 (2006).

14 Chiu, C., Xian, W. & Moss, C. F. Flying in silence: Echolocating bats cease vocalizing to avoid sonar jamming. Proc Natl Acad Sci U S A 105, 13116–13121, doi:10.1073/pnas.0804408105 (2008).

15 von der Emde, G. & Schwarz, S. Imaging of objects through active electrolocation in Gnathonemus petersii. J Physiol Paris 96, 431–444, doi:10.1016/S0928-4257(03)00021-4 (2002).

16 Lissmann, H. W. & Machin, K. E. The mechanism of object location in _Gymnarchus niloticus_ and similar fish. J. Exp. Biol 35, 451–486 (1958).

17 Carlson, B. A. & Gallant, J. R. From sequence to spike to spark: evo-devo-neuroethology of electric communication in mormyrid fishes. Journal of neurogenetics 27, 106–129 (2013).

18 Hopkins, C. D. Neuroethology of electric communication. Annual review of neuroscience 11, 497–535 (1988).

19 Hopkins, C. D. in Electroreception (eds T.H. Bullock & W. Heiligenberg) 527–571 (John Wiley & Sons, Inc., 1986).

20 Moller, P. Electric signals and schooling behavior in a weakly electric fish, _Marcusenius cyprinoides_ L. (Mormyriformes). Science 193, 697–699 (1976).

21 Moller, P., Serrier, J., Belbenoit, P. & Push, S. Notes on ethology and ecology of the Swashi River mormyrids (Lake Kainji, Nigeria). Behavioral Ecology and Sociobiology 4, 357–368 (1979).

22 Gebhardt, K., Alt, W. & von der Emde, G. Electric discharge patterns in group-living weakly electric fish, Mormyrus rume (Mormyridae, Teleostei). Behaviour, 623-644 (2012).

23 Arnegard, M. E. & Carlson, B. A. Electric organ discharge patterns during group hunting by a mormyrid fish. Proc Biol Sci 272, 1305–1314, doi:10.1098/rspb.2005.3101 (2005).

24 Russell, C. J., Myers, J. P. & Bell, C. C. The echo response in Gnathonemus petersii. J. Comp. Physiol 92, 181–200 (1974).

25 Kramer, B. Electric organ discharge interaction during interspecific agonistic behaviour in freely swimming mormyrid fish. A method to evaluate two or more. J. Comp. Physiol 93, 203–236 (1974).

26 Rother, D. et al. Electric images of two low resistance objects in weakly electric fish. BioSystems 71, 169–177 (2003).

27 Bell, C. C. in Electroreception (eds T.H. Bullock & W. Heiligenberg) 423–452 (John Wiley & Sons, Inc., 1986).

28 Fortune, E. S. The decoding of electrosensory systems. Current opinion in neurobiology 16, 474–480 (2006).

29 Push, S. & Moller, P. Spatial aspects of electrolocation in the mormyrid fish, _Gnathonemus petersii_. J. Physiol. (Paris) 75, 355–357 (1979).

30 Pedraja, F., Hofmann, V., Goulet, J. & Engelmann, J. Task-related sensorimotor adjustments increase the sensory range in electrolocation. Journal of Neuroscience 40, 1097–1109 (2020).

31 Babineau, D., Longtin, A. & Lewis, J. E. Modeling the electric field of weakly electric fish. J Exp Biol 209, 3636–3651, doi:10.1242/jeb.02403 (2006).

32 Chen, L., House, J. L., Krahe, R. & Nelson, M. E. Modeling signal and background components of electrosensory scenes. J Comp Physiol A Neuroethol. Sens. Neural Behav. Physiol 191, 331–345 (2005).

33 Post, N. & von der Emde, G. The ‘novelty respons’ in an electric fish: response properties and habituation. Physiol. Behav 68, 115–128 (1999).

34 Enikolopov, A. G., Abbott, L. F. & Sawtell, N. B. Internally Generated Predictions Enhance Neural and Behavioral Detection of Sensory Stimuli in an Electric Fish. Neuron 99, 135–146 e133, doi:10.1016/j.neuron.2018.06.006 (2018).

35 Worm, M. et al. Evidence for mutual allocation of social attention through interactive signaling in a mormyrid weakly electric fish. Proceedings of the National Academy of Sciences 115, 6852–6857 (2018).

36 Moller, P. Electric Fishes: History and Behavior. (Chapman and Hall, 1995).

37 von der Emde, G. Discrimination of objects through electrolocation in the weakly electric fish, _Gnathonemus petersii_. J. Comp. Physiol. A 167, 413–421 (1990).

38 Hall, J. C., Bell, C. & Zelick, R. Behavioral evidence of a latency code for stimulus intensity in mormyrid electric fish. J. Comp Physiol. A, 29-39 (1995).

39 Bell, C. C. Mormyromast electroreceptor organs and their afferents in mormyrid electric fish: III. Physiological differences between two morphological types of fibers. J. Neurophysiol 63, 319–332 (1990).

40 Toerring, M. J. & Moller, P. Locomotor and electric displays associated with electrolocation during exploratory behavior in mormyrid fish. Behav. Brain Res 12, 291–306 (1984).

41 Russell, C. J. & Bell, C. C. Neuronal responses to electrosensory input in the mormyrid valvula cerebelli. J. Neurophysiol 41, 1495–1510 (1978).

42 Sawtell, N. B., Mohr, C. & Bell, C. C. Recurrent feedback in the mormyrid electrosensory system: cells of the preeminential and lateral toral nuclei. J. Neurophysiol 93, 2090–2103 (2005).

43 Prechtl, J. C. et al. Sensory processing in the pallium of a mormyrid fish. J. Neurosci 18, 7381–7393 (1998).

44 Berdahl, A., Torney, C. J., Ioannou, C. C., Faria, J. J. & Couzin, I. D. Emergent sensing of complex environments by mobile animal groups. Science 339, 574–576 (2013).

45 Worm, M., Kirschbaum, F. & von der Emde, G. Social interactions between live and artificial weakly electric fish: Electrocommunication and locomotor behavior of Mormyrus rume proboscirostris towards a mobile dummy fish. PLoS One 12, e0184622 (2017).

46 Schuster, S. Count and spark? the echo response of the weakly electric fish gnathonemus petersii to series of pulses. J. Exp. Biol 204, 1401–1412 (2001).

47 Bell, C. C., Han, V. & Sawtell, N. B. Cerebellum-like structures and their implications for cerebellar function. Annu. Rev. Neurosci 31, 1–24 (2008).

48 Nieuwenhuys, R. & Nicholson, C. in Neurobiology of Cerebellar Evolution and Development (ed R. Llinas) 107–134 (Am. Med. Assoc, 1969).

49 Sukhum, K. V., Shen, J. & Carlson, B. A. Extreme enlargement of the cerebellum in a clade of teleost fishes that evolved a novel active sensory system. Current Biology 28, 3857-3863. e3853 (2018).

50 Finger, T. E., Bell, C. C. & Russell, C. J. Electrosensory pathways to the valvula cerebelli in mormyrid fish. Exp. Brain Res 42, 23–33 (1981).

## Methods References

51 Bell, C. C., Myers, J. P. & Russell, C. J. Electric Organ Discharge Patterns during Dominance Related Behavioral Displays in Gnathonemus-Petersii(Mormyridae). J Comp Physiol 92, 201–228, doi:Doi 10.1007/Bf00694506 (1974).

52 Mathis, A. et al. DeepLabCut: markerless pose estimation of user-defined body parts with deep learning. Nat Neurosci 21, 1281–1289, doi:10.1038/s41593-018-0209-y (2018).

53 Arthur, D. & Vassilvitskii, S. in Proceedings of the eighteenth annual ACM-SIAM symposium on Discrete algorithms. 1027–1035.

54 Braun, E., Geurten, B. & Egelhaaf, M. Identifying prototypical components in behaviour using clustering algorithms. PloS one 5, e9361 (2010).

55 Migliaro, A., Caputi, A. A. & Budelli, R. Theoretical analysis of pre-receptor image conditioning in weakly electric fish. PLoS. Comput. Biol 1, 123–131 (2005).

56 Engelmann, J., Bacelo, J., van den, B. E. & Grant, K. Sensory and motor effects of etomidate anesthesia. J Neurophysiol 95, 1231–1243 (2006).

57 Requarth, T. & Sawtell, N. B. Plastic corollary discharge predicts sensory consequences of movements in a cerebellum-like circuit. Neuron 82, 896–907 (2014).

58 Bell, C. C., Grant, K. & Serrier, J. Corollary discharge effects and sensory processing in the mormyrid electrosensory lobe: I. Field potentials and cellular activity in associated structures. J. Neurophysiol 68, 843–858 (1992).

59 Bell, C. C., Caputi, A. & Grant, K. Physiology and plasticity of morphologically identified cells in the mormyrid electrosensory lobe. J. Neurosci 17, 6409–6422 (1997).

60 Mohr, C., Roberts, P. D. & Bell, C. C. Cells of the mormyromast region of the mormyrid electrosensory lobe: I. Responses to the electric organ corollary discharge and to electrosensory stimuli. J. Neurophysiol 90, 1193–1210 (2003).

61 Kobak, D. et al. Demixed principal component analysis of neural population data. elife 5, e10989 (2016).

