## Supplemental for "Collective Sensing in Electric Fish"

**Extended Data**

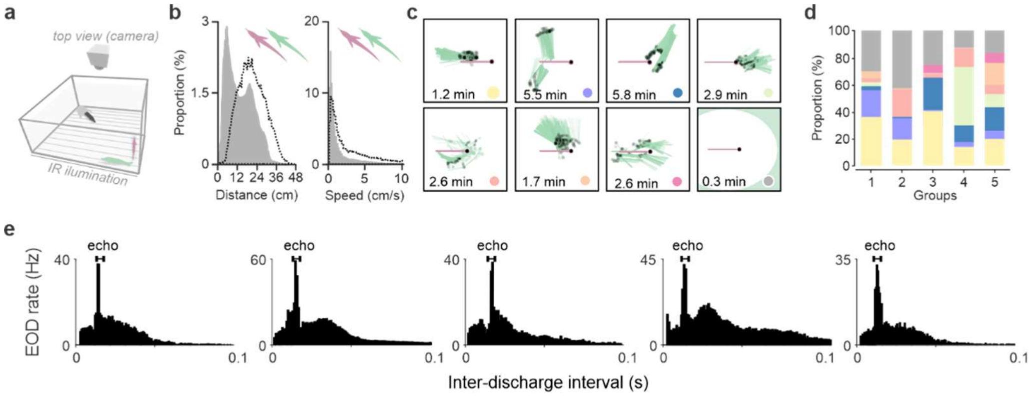


**Extended Data Fig. 1: Stable group behavior in *Gnathonemus petersii*. a,** Experimental setup for video tracking of groups consisting of one large (black) and two smaller (green and purple) *Gnathonemus petersii*. **b,** Distance between the two smaller fish and their average speed during the day (gray) and night (dashed line) (n=5 groups). **c,** Stable daytime spatial configurations between two smaller fish identified using k-means cluster analysis (n=5 groups of 3 fish). A third larger fish was also present and occupied a shelter located in the corner of the tank. For each cluster, the positions of the two fish (lines) are shown for 1000 randomly selected video frames. The heads of the fish are indicated by black dots. The number indicates the average continuous time fish spent in each configuration. **d**, Closely-spaced configurations occupied the majority of the light-cycle in all 5 groups of fish tested (colors). Gray indicates periods when fish were >15 cm apart. **b,** EOD interval histograms during daytime for each group. The prominent peak at approximately 12 ms latency is known as the echo response.

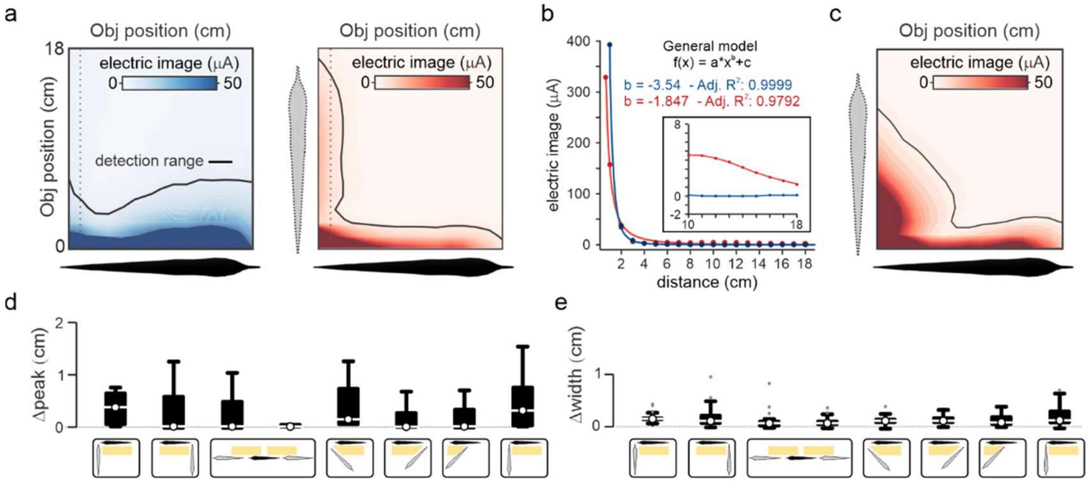


**Extended Data Fig. 2: Physical characteristics of conspecific electrical images.** **a**, Maps of the maximal electric image generated by the fish's own EOD (left) and the EOD of a conspecific in a right-angle configuration (right) for a 1 cm metal sphere placed at various distances and locations along the length of the fish. Black line indicates the range of object detection based on prior behavioral studies. **b**, Maximal electrical images as a function of distance (along the dashed line in **c**) for a 1 cm metal sphere with power law fits to the data. Self-images (blue), Cons-images (red) Inset shows a magnification between 10-18 cm. **c**, Same as in **a**, but with the internal conductivity of the model conspecific (dashed outline) set equal to water in order to visualize the 'funneling' effects of the conspecific fish's body on electrical images. **d,** Difference in electric image peak location on the skin for self- and cons-images for a range of object positions (yellow) for eight different conspecific spatial configurations. Related to **Fig. 1g,h**. **e,** Same as in **d** for electrical image width. Related to **Fig. 1g,i**

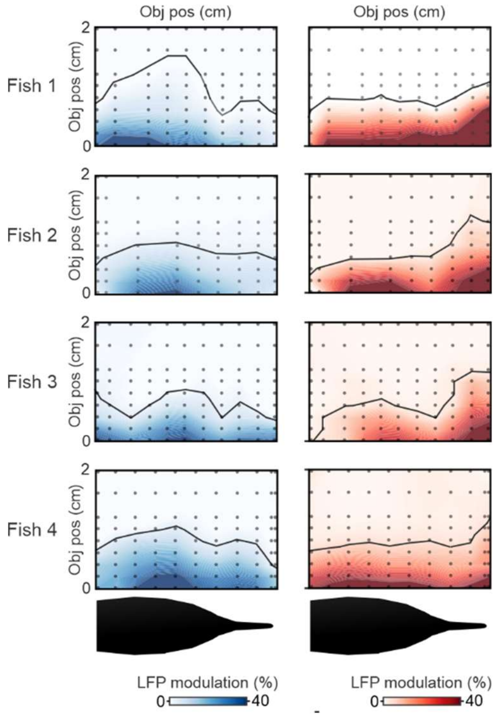


**Extended Data Fig. 3: Neural correlates for electrolocation range extension by cons-EODs.** Maps of LFP amplitude modulation evoked by a plastic object by a 2 cm plastic rod placed at different locations near the fish (black dots). Columns are separate recording locations on the ELL map matched to the rostro-caudal locations of the object. Black lines indicate a 5% modulation of LFP amplitude. For locations near the chin appendage (left), LFP amplitude was modulated by objects at further distances for cons- (red) versus self-EODs (blue). Related to **Fig. 2d,f**.

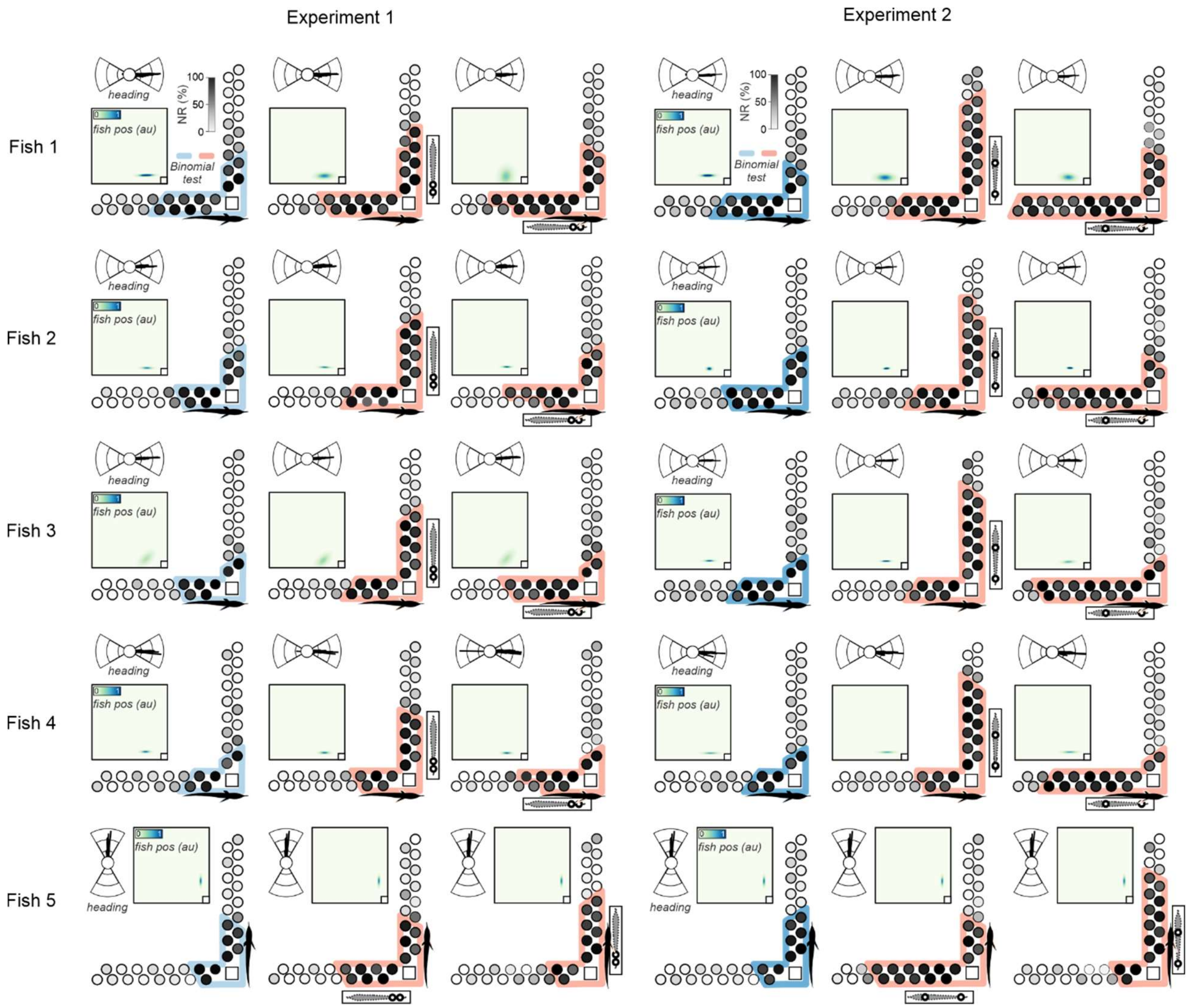


**Extended Data Fig. 4: Behavioral evidence for enhanced electrolocation range based on exogenous EODs.** Each set of three L-shaped plots shows NR probability for each object (grayscale) with mimic pulses off (blue; left) versus on (red; middle, right). Colored contours indicate behavioral detection range (Binomial test: $p < 0.05$). Insets: top, histogram of the angular orientation of the fish for each condition across the entire ~7 days experiment. Bottom, heatmap of the distribution of fish positions in the experimental tank for each condition across the entire ~7 days experiment. In *Experiment 2* the spacing between poles of the EOD mimic was increased, simulating a larger fish.

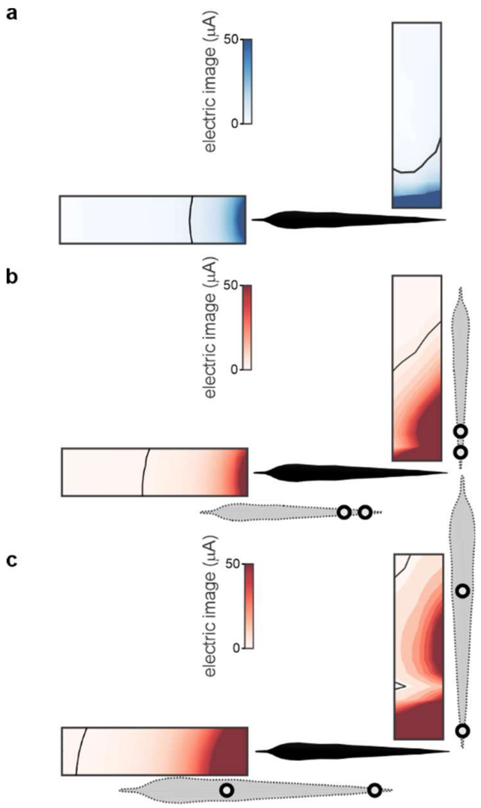


**Extended Data Fig. 5: Direction and extent of behavioral range extension matches boundary element model predictions. a**, Boundary element simulations of maximal electrical images for a 1 cm metal sphere generated by the fish's own EOD. **b**, Same as in **a** but for cons-images generated by an EOD mimic with 4 cm spacing between electrodes used in *experiment 1*. Compare to behavioral results in Fig. **3d**. **c**, Same as in **a** but for cons-images generated by an EOD mimic with 16 cm spacing between electrodes used in *experiment 2* to simulate a larger conspecific. Compare to behavioral results in **Fig. 3e**.

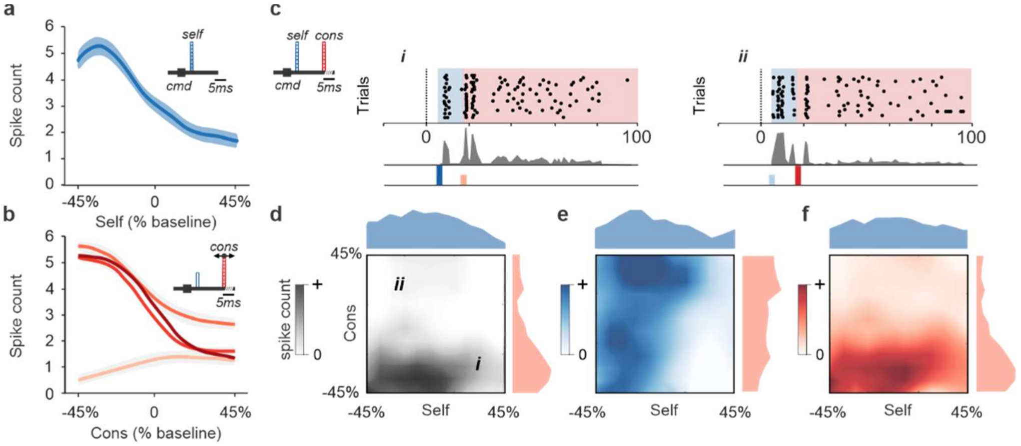


**Extended Data Fig. 6. Concurrent processing of self- and cons-EODs in ELL output cells.** **a,** Response of an I-type ELL output cell to modulations of the amplitude (3% steps) of a self-EOD pulse (i.e. an artificial pulse locked to the fish's spontaneously emitted EOD motor command at a 4.5 ms delay corresponding to the naturally occurring EOD). **b**, Response of the same cell to modulations of the amplitude of cons-EOD pulses delivered across a range of delays. **c,** Response of an example I-type ELL output cell to independent modulations of self- and cons-pulses (12 ms delay). Rasters and peri-stimulus time histograms illustrating responses to self- (blue) and cons-pulses (red) of different amplitudes (i and ii). Shaded rectangles indicate time windows used for quantifying responses to self- (blue) and cons-pulses (red). **d**, Same unit as in **c**. Grayscale heatmap indicates the total spike count in 100 ms window following the EOD motor command. **e,f,** Responses of the same unit calculated in separate time windows following self- and cons-pulses.

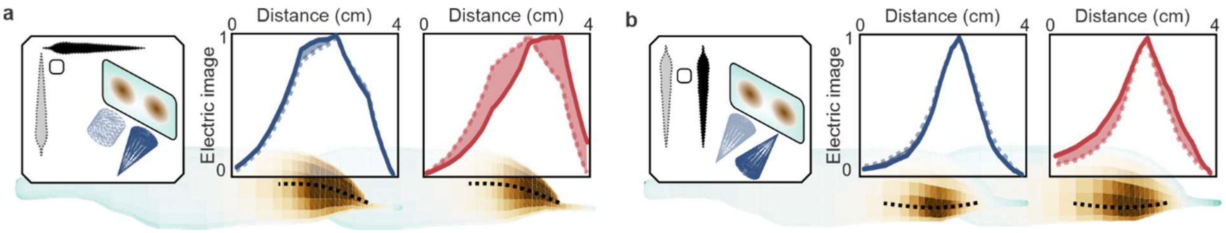


**Extended Data Fig. 7: Self- and cons-electrical images provide different views of the same scene. a**, Boundary element model simulations of self- (blue) and cons- (red) electrical images of a metal cylinder (dashed) and a metal cone (solid) both 1 cm in diameter. Normalized electrical images are shown on a 3D model of the fish (bottom) and as 2D profiles (above). Black dashed lines indicate the location of the image profile on the head of the fish. Conspecific fish orientation is indicated in gray. **b**, Same as in **a** but for a different conspecific orientation and for two different orientations of a 1 cm diameter cone. Dashed and solid lines indicate the base of the cone closer and farther from the fish, respectively. In both scenarios, the spatial profiles of cons-images show greater differences (shaded area) for the two object configurations than the self-images, implying that cons-images could enhance object discrimination.
